# Days Gained: A Simulation-Based, Response Metric in the Assessment of Glioblastoma

**DOI:** 10.1101/209056

**Authors:** Gustavo De Leon, Kyle W. Singleton, Kristin R. Swanson

## Abstract

We show the application of a minimally based, patient-specific mathematical model in the evaluation of glioblastoma response to therapy. Days Gained uses computational models of glioblastoma growth dynamics derived from clinically acquired magnetic resonance imaging (MRI) to compare the post-treatment tumor lesion to the *expected* untreated tumor lesion at the same time point. It accounts for the inter-patient variability in growth dynamics and response to therapy. This allows for the accurate assessment of therapeutic response and provides insight into overall survival as it relates to treatment response.

## I. Assessment of Therapeutic Response in Glioblastoma

Glioblastoma (GBM) is an aggressive primary brain cancer associated with a median survival of 7-8 months (untreated) and 12-14 months (treated) [1]. This short survival is mainly attributed to the aggressive and infiltrative properties of GBM, lack of effective therapeutic interventions, and limited ability to identify early treatment response predictive of survival or other outcomes. While new knowledge continues to elucidate the underlying kinetics of GBM growth, therapeutic decisions continue to rely on ‘one-size-fits-all’ approaches rather than ‘targeted’ or ‘patient-specific’ models. Updated response assessments can help avoid premature discontinuation of potentially beneficial therapy and provide guidance for individualized therapeutic schedules.

Currently, the Response Assessment in Neuro-oncology (RANO) criteria serves as the standard clinical guideline in evaluating therapeutic response for patients with GBM [2]. The assessment is conducted via MRI and their respective sequences. The T1-weighted with gadolinium contrast enhancement (T1Gd) MRI sequence is used to identify the tumor burden associated with a higher density of tumor cells. The T2 MRI sequence highlights the edema associated with the distribution of tumor cells of lower density. The RANO criteria compares pre- and post-treatment tumor burden but does not account for differential growth kinetics across tumors. Even when combined with patient’s clinical symptoms, decisions’ pertaining to changes in treatment measurements from two pre-treatment MRIs are required to run the UVC and estimate an untreated tumor burden at post-treatment time points. The UVC can then be directly compared against actual tumor lesions measured posttreatment to compute a Days Gained response value (*Fig. 1*).

**Figure 1.**
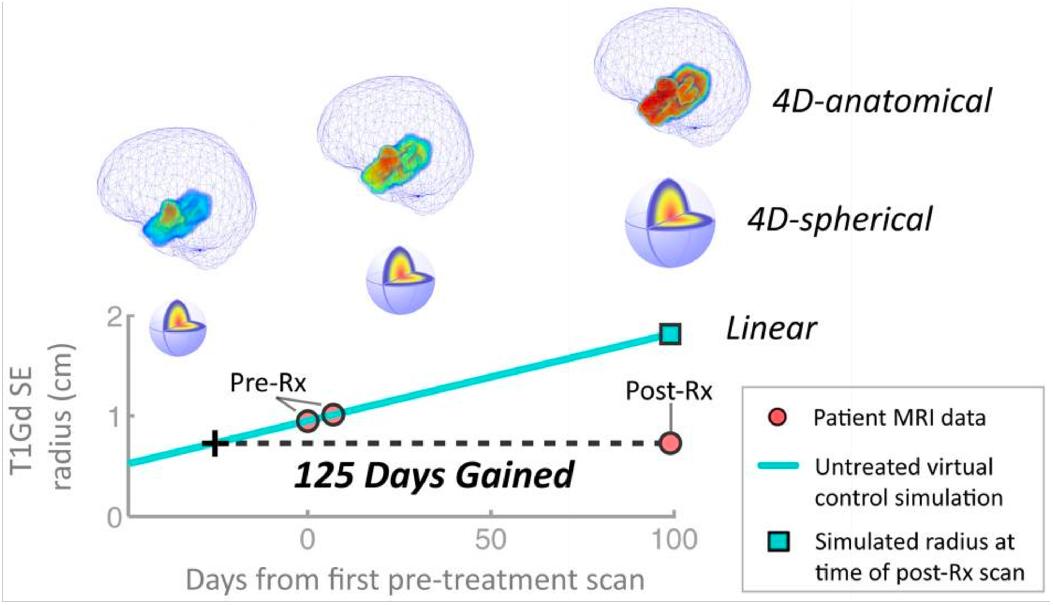
Representation of the three models used as untreated virtual controls in order to compute Days Gained scores from patient-specific MRI data. The T1Gd spherically-equivalent (SE) radius is derived from volumetric extractions of pre-treatment MRI scans to generate a UVC and compare it to the tumor lesion at a post-treatment time point.

## II. Discriminatory Power of Days Gained

DG has been validated in two investigations of patients receiving radiation therapy, demonstrating a significant discrimination between patients in progression-free survival and overall survival. The first investigation included 33-newly diagnosed GBM patients receiving first-line radiotherapy [3]. The second investigation (*Fig. 2*) expanded the cohort to 63 newly diagnosed GBM patients [4]. In addition, the second study compared multiple methods for estimating the tumor burden of UVC models, determining that each version of the model revealed comparable results and a simplified linear model could provide an efficient and accurate computation of a UVC.

**Figure 2.**
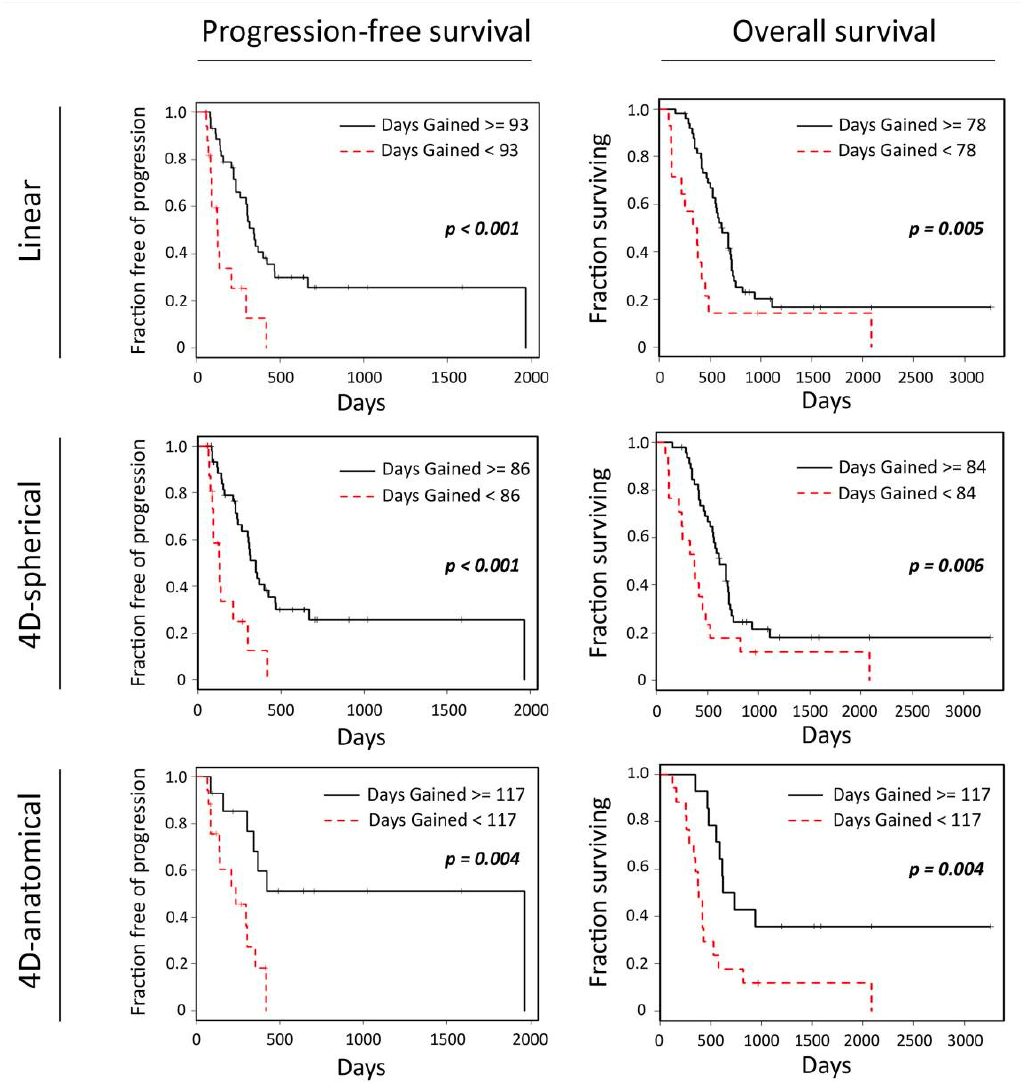
Kaplan-Meier analyses comparing the PFS and OS of patients classified according to Days Gained score. This study incorporated three models differing in level of complexity (top = low complexity, middle = medium complexity, bottom = high complexity). DG scores computed from the least complex, linear model demonstrates a level of significant discrimination similar to the more complex, 4D-anatomical and 4-D spherical models.

## III. Computing Days Gained Scores

There are three models used to simulate UVCs representative of tumor growth at distinct levels of detail. Each of these 4D UVC simulations are generated on the basis of a Proliferation-Invasion (PI) model that accounts for two patient-specific parameters: tumor’s net diffusive capacity (D) and tumor’s net proliferation rate (p) [5-12]. The 4D-anatomical model simulates tumor growth based on a reaction-diffusion differential equation parameterized by D and ρ. The diffusion values of simulated cells take into account the patient’s neuroanatomy using segmented gray matter, white matter, and cerebrospinal fluid regions to dynamically influence the geometric shape of the tumor. The 4D-spherical model simplifies the anatomical model by simulating tumor growth as an expanding sphere without imposing restrictions based on neuroanatomy. Lastly, the linear model simplifies further, assuming a linear radial expansion of the tumor lesion without simulating individual cells. The 4D-anatomical model simulates a growing tumor with the highest accuracy. However, we have shown that the minimal linear model can significantly discriminate patient outcomes in progression free survival and overall survival to the same degree as the complex models. We therefore focus on the application of the linear model since it requires the minimal amount of patient-specific information acquired with routine MRIs and can be quickly applied in the clinical setting.

### A. Linear Model

The linear model simulates the radial expansion of lesions from a given MRI contrast (e.g., T1Gd and T2). Tumor growth is assumed to follow a spherical distribution with initial tumor size estimated as a spherically equivalent (SE) radius, which is derived from segmented tumor volumes visualized in the MRI sequences.

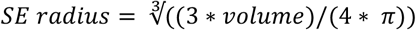

The linear model can be used to estimate predicted SE tumor radii, *y*, at a future time as defined below.

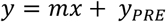

In the linear model, *y* represents the predicted SE radii on T1Gd or T2 imaging, *m* is the pre-treatment growth velocity measured on T1Gd or T2 imaging, *x* refers to the amount of time between the final pre-treatment scan and next time point of interest (typically the first post-treatment scan), and *y_pre_* represents the SE tumor radius extracted from the last pre-treatment T1Gd or T2 scan. To calculate *m*, we take the volumetric difference between two segmented tumors from two pre-treatment scans and divide it by the amount of time between the scans.

The predicted values from this Untreated Virtual Control (UVC) based on the linear model can be compared to the actual tumor size from future post-treatment time scans to determine the degree of therapeutic response. By solving for x in the linear model, we can estimate the amount of time a given therapy deflected tumor growth.

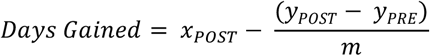

DG scores are calculated by replacing the predicted tumor size, *y*, with the segmented post-treatment T1Gd or T2 SE radius, *y*_POST_. The difference in pre- and post-treatment tumor size is then divide by the tumor growth velocity, m, and subtracted from the amount of time that has passed between scans, *x*_POST_ (in days). DG scores assume a positive tumor growth rate prior to therapeutic intervention. Therefore, these scores are only meaningful for positive values of *m*.

### B. Days Gained in the Clinic

The DG response metric was developed to gain insight in the patient-specific differences that account for the vast heterogeneity among patients, patients’ tumors, and response to therapy. As previously mentioned, the DG metric has shown promise in discriminating patients receiving standard-of-care radiotherapy. In recently analysis, DG scores showed discriminative power in several cohorts receiving different therapies [13, 14]. These therapies include Bevacizumab, an anti-angiogenic drug whose main function is to suppress the development of new blood vessels, and Gamma Knife, a targeted form of radiation therapy with high accuracy.

The application of DG as a response metric is useful in the clinical context of GBM patients receiving different treatments. Using the minimal linear model, DG can be obtained using routinely acquired MRIs without disrupting clinical workflow. Thus, DG scores have the potential to aid physicians in deciding course of treatment or enrolling patients to appropriate clinical trials.

## Acknowledgment

We thank the organizers of the ICBP/PSOC meeting and creators of the on-going Handbook of Mathematical Methods in Cancer Biology and the NIH/NCI for sponsoring the 2017 Mathematical Oncology meeting.

## Notes

We gratefully acknowledge the support of the James S. McDonnell Foundation, the Ivy Foundation, the Mayo Clinic and the NIH (R01 NS060752, R01CA164371, U54CA210180, U54CA143970, U54 CA193489, U01CA220378).

## References

[1] Smoll NR, Schaller K, Gautschi OP. “Long-term survival of patients with glioblastoma multiforme (GBM).” Journal of Clinical Neuroscience. 20.5 (2013): 670–675.

[2] Wen PY, Macdonald DR, Reardon DA, Cloughesy TF, Sorensen AG, Galanis E, DeGroot J, Wick W, Gilbert MR, Lassman AB, Tsien C, Mikkelsen T, Wong ET, Chamberlain MC, Stupp R, Lamborn KR, Vogelbaum MA, van den Bent MJ, Chang SM. “Updated Response Assessment Criteria for High-Grade Gliomas: Response Assessment in Neuro-Oncology Working Group.” Journal of Clinical Oncology. 28.11 (2010): 1963–1972.

[3] Neal ML, et al. “Discriminating Survival Outcomes in Patients with Glioblastoma Using a Simulation-Based, Patient-Specific Response Metric.” Ed. Russell O. Pieper. PLoS ONE. 8.1 (2013): e51951.

[4] Neal ML, et al. “Response Classification Based on a Minimal Model of Glioblastoma Growth Is Prognostic for Clinical Outcomes and Distinguishes Progression from Pseudoprogression.” Cancer Research. 73.10 (2013): 2976–2986.

[5] Swanson KR. (1999). Mathematical modeling of the growth and control of tumors, Dissertation. University of Washington

[6] Swanson KR, Alvord E, Murray J, Rockne R (2010). Method and system for characterizing tumors, United States

[7] Wang CH, Rockhill JK, Mrugala M, Peacock DL, Lai A, et al. “Prognostic significance of growth kinetics in newly diagnosed glioblastomas revealed by combining serial imaging with a novel biomathematical model.” Cancer Research. 69.23 (2009): 9133–9140.

[8] Swanson KR, Rostomily RC, Alvord EC., Jr. “A mathematical modelling tool for predicting survival of individual patients following resection of glioblastoma: a proof of principle.” British Journal of Cancer. 98.1 (2008):113–119.

[9] Harpold HL, Alvord EC Jr, Swanson KR. ‘The evolution of mathematical modeling of glioma proliferation and invasion.” Journal of Neuropathology & Experimental Neurology. 66.1 (2007): 1–9.

[10] Swanson KR, Alvord E, Jr, Murray J. “Virtual resection of gliomas: effect of extent of resection on recurrence.” Mathematical and Computer Modeling. 37.11 (2013): 1177–1190.

[11] Baldock A, Yagle K, Born DE, Ahn S, Trister AD, Neal M, Johnston SK, Bridge CA, Basanta D, Scott J, Malone H, Sonabend AM, Canoll P, Mrugala MM, Rockhill JK, Rockne RC, Swanson KR. “Invasion and proliferation kinetics in enhancing gliomas predict IDH1 mutation status.” Neuro-Oncology. 16.6 (2014): 779–786.

[12] Swanson KR, and Murray JD. K. R. “Dynamics of a model for brain tumors reveals a small window for therapeutic intervention,” Discrete and Continuous Dynamical Systems – Series B. 4.1(2003): 289–295.

[13] Singleton, Kyle et al. “Discrimination of clinically impactful treatment response in recurrent glioblastoma patients receiving bevacizumab treatment.” Abstract. Society of Neuro-Oncology. 2017. In press.

[14] Singleton, Kyle et al. “Role of pre-treatment tumor dynamics and imaging response in discriminating glioblastoma survival following gamma knife.” Abstract. Society of Neuro-Oncology. 2017. In press.

